# Learning dynamic image representations for self-supervised cell cycle annotation

**DOI:** 10.1101/2023.05.30.542796

**Authors:** Kristina Ulicna, Manasi Kelkar, Christopher J Soelistyo, Guillaume T Charras, Alan R Lowe

**Affiliations:** The Alan Turing Institute, London NW1 2DB, United Kingdom; Institute of Structural and Molecular Biology, University College London, London WC1E 6BT, United Kingdom; London Centre for Nanotechnology, University College London, London WC1H 0AH, United Kingdom; Department of Cell and Developmental Biology, University College London, London WC1E 6BT, United Kingdom; Institute for the Physics of Living Systems, University College London, London WC1E 6BT, United Kingdom

**Author notes:** Correspondence to: Kristina Ulicna < >, Alan R Lowe < >.

## Abstract

Time-based comparisons of single-cell trajectories are challenging due to their intrinsic heterogeneity, autonomous decisions, dynamic transitions and unequal lengths. In this paper, we present a self-supervised framework combining an image autoencoder with dynamic time series analysis of latent feature space to represent, compare and annotate cell cycle phases across singlecell trajectories. In our fully data-driven approach, we map similarities between heterogeneous cell tracks and generate statistical representations of single-cell trajectory phase durations, onset and transitions. This work is a first effort to transform a sequence of learned image representations from cell cycle-specific reporters into an unsupervised sequence annotation.

## 1. Introduction & Background

During its lifetime, a cell passes through a series of well-characterised, morphologically distinct cycling stages that prepare it to divide (Fig.1, App.A). Quantitative cell cycle analysis is an essential tool for understanding cell behavior and dynamics of their growth and division. In recent years, multigenerational lineage tracing has enabled measuring cell behavior at the single-cell level over time. However, accurate identification and labeling of cell cycle phases dynamics along trajectories with variable durations remains challenging as it requires meaningful interpretation of low-resolution signals that vary continuously over time (Fig.1).

**Figure 1.**
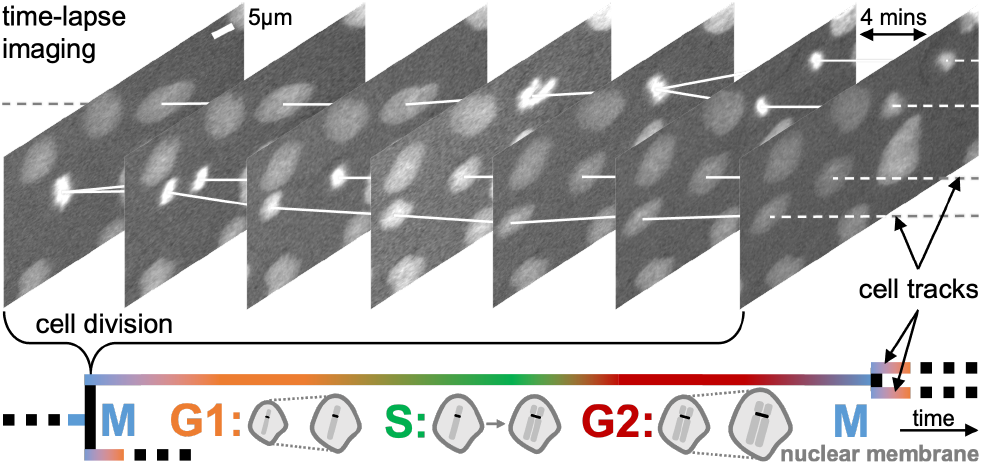
The objective to annotate continuous measurements, such as cell cycle phases, in single-cell trajectories from raw timelapse image data capturing cell cycle-specific reporter dynamics over the <u>G</u>rowth, <u>S</u>ynthesis and <u>M</u>itotic phases of variable lengths.

In this paper, we use the cell cycle phases as an exemplar to present a novel, supervision-free approach for automated representation, comparison and annotation of continuous processes along single-cell trajectories. Our data-driven trajectory analysis is based on (i) self-supervised extraction of latent features from cell cycle-specific image data and (ii) inference of phase labels via parameter-free dynamic time warping (DTW) from a miniature annotated cell library.

Previous analyses addressing the morphodynamic transition challenges have sourced from latent image representations, however, these methods either rely on compressing the entire trajectory with variable durations into single, fixed-length embedding (Wu et al., 2022), or truncate the time-resolved tracks to an identical, pre-defined length (Soelistyo et al., 2022). Conversely, the length-invariant approaches are either driven by hand-crafted features (El-Labban et al., 2014) or yield poorly interpretable continuous latent space clustering (Ji et al., 2017), which complicates discrete categorical cell cycle phase annotation (Rappez et al., 2020).

Our approach contributes to the associated challenges of cell cycle phase labeling three-fold. First, our cell-specific image autoencoder learns compressed latent representations of individual cell images and captures key instantaneous features of a cell cycle-specific reporter. This step addresses the difficulty of quantifying low-resolution signals and compensates for complicated feature hand-crafting. Second, the sequence of image-specific latent space embeddings is employed to localise regions of (dis-)similarity between two trajectories by dynamic matching. We leverage this self-supervised strategy to infer phase labels of an unseen trajectory from an annotated cell library to categorically label cell phases. This automated labeling step allows time-based comparisons in the context of entire trajectories of unequal lengths. Third, extending beyond categorical labeling, our approach enables the quantitative statistical representation of label distribution in cell trajectories for a supervision-free annotation of new data, enriched with an uncertainty measure. We challenge the concept of categorical labeling of continuous processes by proposing a flexible, more appropriate alternative.

## 2. Methods & Workflow

### 2.1. Live-cell imaging & single-cell trajectory dataset

The image dataset was acquired by automated time-lapse microscopy via live-cell fluorescence imaging of Madin–Darby canine kidney (MDCK) epithelial cells (Bove et al., 2017), expressing both *H2b-GFP* (constitutive histone marker for nuclei detections) and *PCNA-iRFP* (cell cycle phase-specific protein with morphological transitions) reporter systems.

The cells were segmented (Weigert et al., 2020), classified (Ulicna et al., 2022) and tracked (Ulicna et al., 2021) to reconstruct multi-generational lineage trees (Fig.1). Singlecell trajectories had their image patches centered at the cell nucleus along the track and cropped at *≈*10*×*10*µm* scale (Fig.2, App.B). The complete, hand-annotated dataset consists of 204 fully-resolved single-cell trajectories from 4 movies, divided into 35 reference *vs*. 169 query cell tracks.

**Figure 2.**
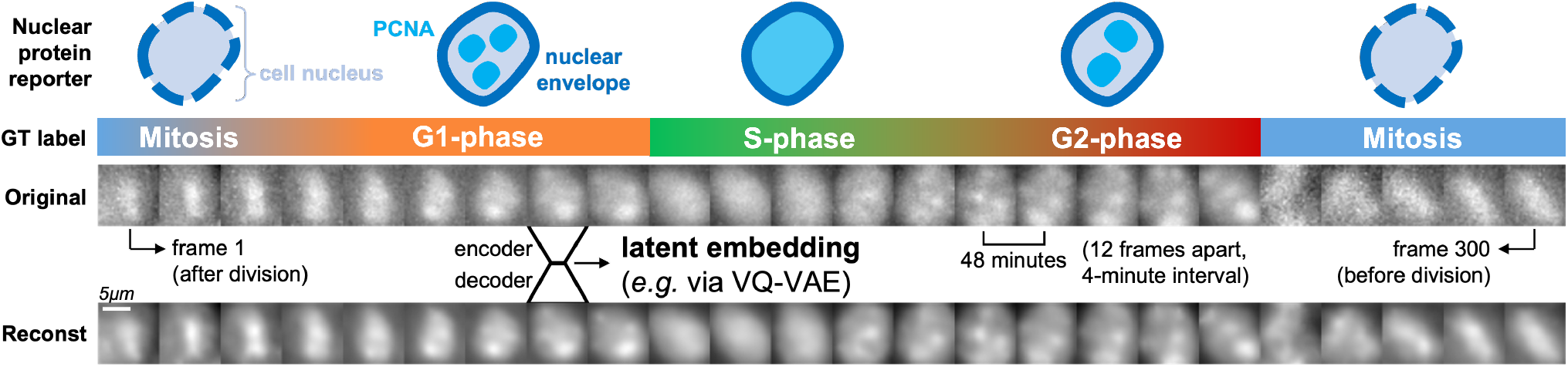
Characteristic PCNA morphology throughout the cell cycle. Hand-annotated phases and transitions along an original single-cell image sequence with corresponding encoded-decoded *VQ-VAE* reconstructions, capturing key image features in form of latent embeddings. These include coarse information, such as cell nucleus size and shape, with detailed patterns of reporter signal distribution.

### 2.2. Latent image representation learning with *VQ-VAE*

The *VQ-VAE* architecture (Oord et al., 2018) is formed of a convolutional *encoder* which compresses an original image *x* into a lower-dimensional representation *z*_*e*_(*x*), which is then *quantized* by a nearest neighbour look-up, or *ℓ*_2_ norm, along the channel dimension to a latent embedding space “codebook” (Fig.3). The quantized discrete embeddings *z*_*q*_(*x*) are then passed through a convolutional *decoder* to produce the reconstruction image *x*^*′*^. All downstream analysis uses the flattened *L*-dimensional representation, where *L∈* ℝ^*h×w×d*^, *i*.*e*. the quantized latent embedding dimensionality. These are extracted per-image along the entire trajectory order of *T* time points in length (Fig.3, App.B).

**Figure 3.**
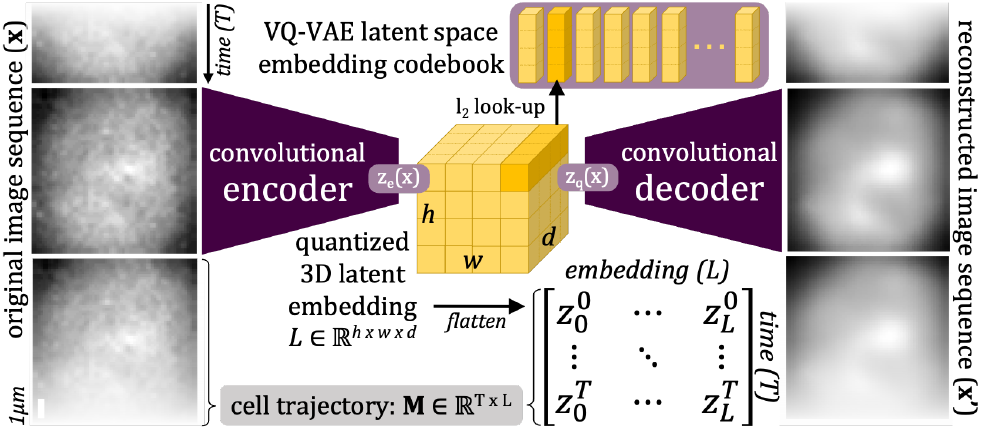
Latent representation learning. *VQ-VAE* is trained to learn a low-dimensional image representation from single-cell image patches. The entire trajectory is represented as *M ∈* ℝ^*T×L*^.

### 2.3. Dynamic time warping of trajectory latent features

Latent feature embeddings are extracted from a pair of arbitrarily long reference and query image sequences to compute a dynamic warping matrix using *ℓ*_2_ distance (Fig.4). The warping matrix is traversed by backward algorithm (Berndt & Clifford, 1994) to identify and locate the minimum-cost alignment path used to infer trajectory phase annotations from the hand-annotated reference cell. Guided by the alignment path, each unannotated query cell position is mapped to the reference cell position to which it warps closest and copies the corresponding reference label (Fig.4, App.C).

**Figure 4.**
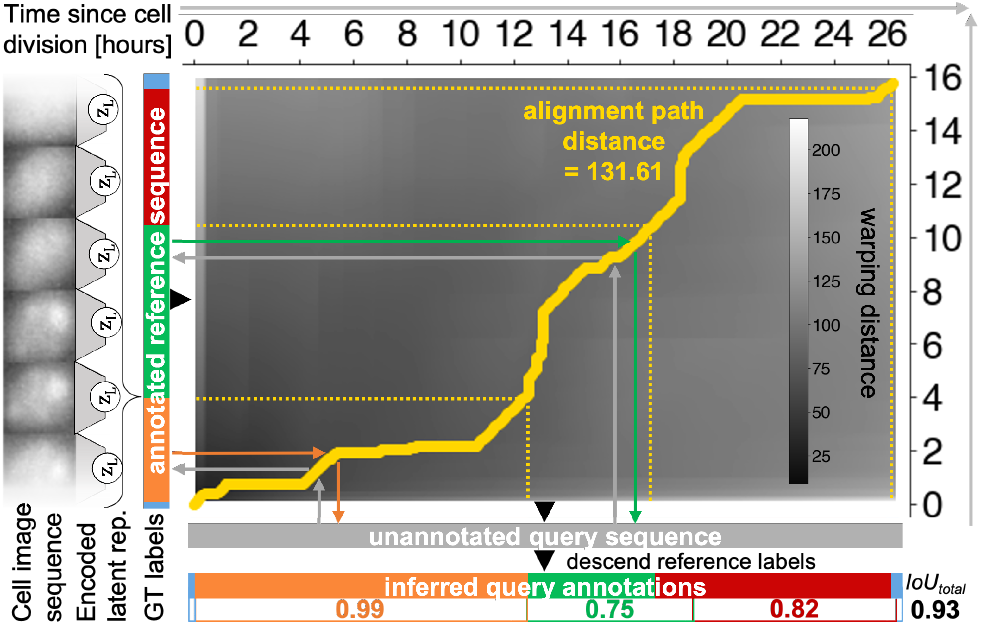
Supervision-free reference-guided labelling of query trajectories with dynamic time warping of latent representations. The *VQ-VAE*-encoded latent embeddings (left) from both reference and query image sequences are warped to compute their best alignment path (thick yellow line) from the warping matrix. Query phase labels (horizontal bars) are inferred from the path (arrows) and follow the categorical labels from a hand-annotated reference (vertical bar). Scores reflect overlap with GT annotation.

### 2.4. Evaluation of categorical phase annotation fidelity

Each “inferred” categorical annotation (from single reference or most common overlap across *N* references, Fig.5) is contrasted to hand-annotated ground truth (GT) via intersection <u>o</u>ver <u>u</u>nion score as 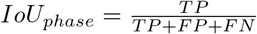, where the *TP* (true +) are instances where the position’s predicted label matches GT, whilst the *FP* (false +) covers instances where predicted label is not in GT, and the *FN* (false −) covers instances where label is in GT but misses in prediction. The overall 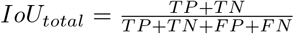 score reflects the ratio of correct detections (true + and −) to all detections (*i*.*e*. sequence agreement or accuracy, Fig.5).

**Figure 5.**
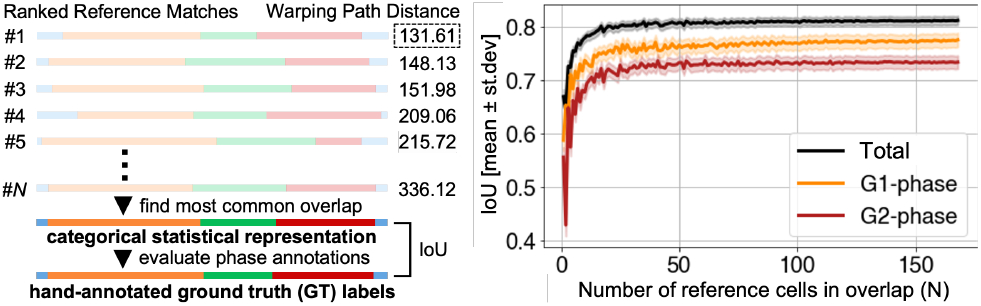
Improving annotation fidelity by considering *N* reference sequences and finding most common overlap per each trajectory position. Such “categorical statistical representation” is evaluated against GT hand-annotations (left). IoU scores (35 queries) initially improve but saturate at *N ≈* 20 references (right).

## 3. Experiments & Results

### 3.1. Motivation behind individual steps of the workflow

This section deploys image latent encodings to perform dynamic time warping between two sequences, allowing the cell cycle phase annotation. Importantly, we aim to highlight the versatility of our workflow which is agnostic to the choice of a feature extractor (§3.2), including pre-trained “off-the-shelf” models or hand-crafted features. Although model explainability is beyond the scope of this work, we implemented an image autoencoder (§2.2) with a motivation to learn *meaningful* latent embeddings which can be verified through image reconstructions (Fig.2). Additionally, custom-trained models allow for latent space dimensionality control, which can be optimised and/or experimented with.

The DTW approach choice (§2.3) could also be substituted by simpler supervision-free methods, such as search for closest image embedding from the reference library on per-image basis without considerations of the time flow. This strategy, however, yields noisy annotations with frequent back-and-forth flipping between states. Such noise, concentrated at phase transitions, is difficult to clean and disables reliable sequence partitioning into cell cycle phase regions.

Approaching the problem with a hidden Markov model (HMM) also poses conceptual limitations as the *Markov property* dictates that the future state depends only on the present state and not on past history. A HMM represents transitions purely statistically, with the state changes guided by (known) transition probabilities. Not only does this homogenise the variable phase lengths in trajectories, but calculating such probabilities requires input of prior knowledge of the modelled system. Unlike HMMs, the supervision-free DTW finds structure by establishing a memory of cell cycle event order as it monitors event flow over time in the context of whole trajectory, which extends its utility to cases where phase durations are variable (§3.2).

### 3.2. Phase annotation with real & distorted trajectories

To evaluate our supervision-free annotation methodology, we initially focused on track instances where the overall length of the two trajectories is different and the relative duration of the individual cell cycle phases to the whole trajectory also differs. Using a representative trajectory pair, we confirm the high performance of our strategy (Fig.4).

To increase the representational power of each annotation to more references, we inferred the phase labels for 35 different query cells by warping each cell to 169 references. To evaluate how many references are needed to provide reliable annotation, we sampled *N* random references and generated a categorical statistical representation (Fig.5). Initially, adding more references significantly improves the fidelity of cell phase labeling, but eventually reaches a point where annotations are not improved nor worsened. This “saturation point” appears with as little as 20 references for overall and phase-specific scores (Fig.5). Whilst recognising that other less reproducible and more stochastic processes could express higher dependency on data quantity, this result hints towards the potential scalability of our approach with larger reference libraries, and to implementations in other time-resolved systems where labeled data availability is sparse.

To extend our technology’s potential to annotate other dynamic processes in cell trajectories, we tested the importance of our cell-specific *VQ-VAE* feature extractor. To benchmark its performance, we replaced the *VQ-VAE* encoder with a custom-trained *β-VAE* model (Higgins et al., 2017) and an IMAGENET pre-trained RESNET-18 network (He et al., 2015) (App.B). Annotating 169 unseen trajectories with inferred overlaps across 35 reference cells, we reveal that our *VQ-VAE*-extracted embeddings yield high annotation fidelity but are interchangeable with other image feature extractors, confirming the versatility of our approach (Table 1). Conversely, warping the trajectories based on hand-crafted features, *e*.*g*. image intensity or region properties, suffers in performance (data not shown), highlighting the importance of deep feature extractors in this task.

**Table 1.**
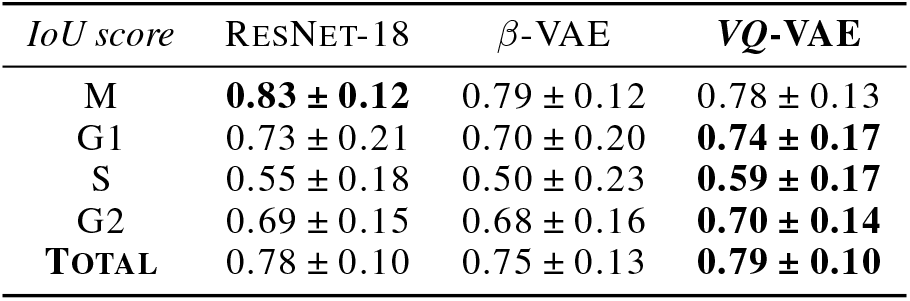
Phase labelling accuracy of 169 query cells warped against 35 reference cells annotated via most common overlap strategy with image latent embeddings from various deep feature extractors.

| $IoU$ score | RESNET-18 | β-VAE | VQ-VAE |
| --- | --- | --- | --- |
| M | **$0.83 \pm 0.12$** | $0.79 \pm 0.12$ | $0.78 \pm 0.13$ |
| G1 | $0.73 \pm 0.21$ | $0.70 \pm 0.20$ | **$0.74 \pm 0.17$** |
| S | $0.55 \pm 0.18$ | $0.50 \pm 0.23$ | **$0.59 \pm 0.17$** |
| G2 | $0.69 \pm 0.15$ | $0.68 \pm 0.16$ | **$0.70 \pm 0.14$** |
| **TOTAL** | **$0.78 \pm 0.10$** | $0.75 \pm 0.13$ | **$0.79 \pm 0.10$** |

Next, we investigated if our time-series matching could be extended to cases where phase durations deviate from expected distributions, such as under pharmacological perturbations which result in evident shortening or prolonging of certain cell cycle phases. To overcome the shortage of real trajectories with out-of-distribution phase durations in our dataset, we synthesized a distorted sequence from a real cell trajectory (Fig.6), and contrasted both variants’ warping pattern against the same trajectory. The dynamic nature of the warping successfully matches the phases in both cases (Fig.6), extending the potential utility of our methodology to cell cycle-targeting pharmacological manipulations.

**Figure 6.**
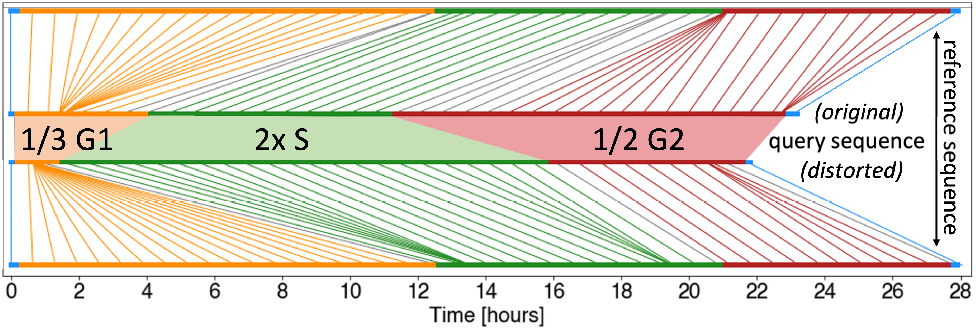
Warping performance on out-of-distribution trajectories. The same reference cell (top and bottom) is warped (vertical lines, gray if labels mismatch) to two sequences: an original (center top) and distorted (center bottom) trajectory, created from every 3^rd^ G1-and every 2^nd^ G2- and by duplicating every S-phase frame.

### 3.3. Statistical representation of cell phase transitions

The continuous nature of PCNA features complicates the labelling of the cell cycle into discrete categories upon both manual and automated annotation. This is most evident at the S/G2 transition which represents a gradual process, unlike at the relatively sudden G1/S switch-like change (Fig.2). Importantly, our approach recapitulates the human annotators’ (un-)certainty when partitioning the trajectory phase regions and can be leveraged to quantify the “label confidence” score along the entire trajectory (Fig.7). Organizing and visualizing the phase annotations of a single query cell via warping to 169 reference cells illustrates where phases hold a strong majority vote *vs*. regions of phase ambiguity, as well as the continuous transitions between them (Fig.7).

**Figure 7.**
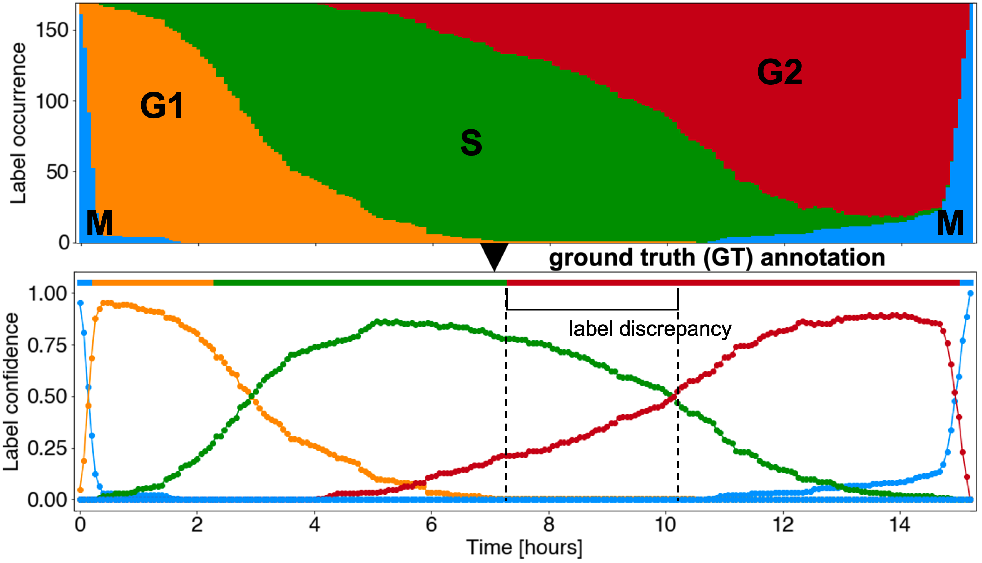
Statistical representation of cell cycle phase annotations. Representative sequence label occurrence per time point (x-axis) inferred from 169 references (y-axis). Visualisation of label confidence, *i*.*e*. the fraction of label’s predicted times, confirms the fast onset of the relatively sharp G1/S transition, whilst the S/G2 transition occurs more gradually (less defined both intrinsically and though PCNA or other reporters), causing a discrepancy (*≈*2.5 hours) between the statistically inferred labels and the GT.

To express the label confidence with high representational power, we compute the frequency of each label, *e*.*g*. the 1^*st*^ trajectory time point has the M-phase inferred 161 out of 169 times (95%, Fig.7). The illustration of this statistical representation can be interpreted as a new, refined categorical annotation, which allows us to express numerical confidence scores to highlight transitional regions. Notably, the per-position label confidence of the cell cycle remains above 50% throughout the trajectory, which is twice as high as pseudo-random label choice, *i*.*e*. 25% if selecting 1 out of 4 labels completely randomly. In general, the highest confidence labels match the GT annotation, except for the ambiguous S/G2 transition which is also difficult to hand-annotate (Fig.7). This motivates us to challenge the consensus of categorical phase annotation and raise the importance of quantifying the certainty of annotating continuous events.

## 4. Discussion & Conclusions

This paper proposes a self-supervised, parameter-free generalizable framework to dynamically find similarities in cell trajectories based on latent image features. Without incorporation of prior knowledge about typical cell cycle-specific characteristics, the cell cycle phase dynamics serve as an exemplar to quantitatively characterize the continuous nature of single-cell trajectory transitions and their rigid labeling.

Certain open questions worth investigating include interpretability of the latent features, *e*.*g*. via ablations or pruning to reveal and evaluate the salient features required for correct annotation. Our framework, currently bound by need for few defined categorical cell labels, enables adaptation to other track annotation purposes in a fully data-driven way.

Despite its limitations, we envisage the applicability of this proposed technology in elucidating the effects of pharmacological perturbations (§3.2) or measuring the dynamics tied to other heterogeneous datasets, *e*.*g*. cell fate determination, signalling pathways, morphogen gradients, *etc*. (Soelistyo et al., 2022; Shakarchy et al., 2023; Gallusser et al., 2023), in biology and beyond.

## Author Contributions

GC and ARL conceived and designed the research. MK performed the microscopy experiments. KU developed and performed the computational analysis. CJS provided computational tools and technical feedback on the manuscript. KU, GC, and ARL evaluated the results and wrote the paper.

## Project Funding

KU and CJS were supported by a BBSRC LIDo studentships. MK was supported by BBSRC grant BB/S009329/1 to ARL and GC.

## A. Appendix The biology of the cell cycle

### Cell cycle as a biological phenomenon

In unicellular as well as multicellular eukaryotes, the **cell cycle** is a ubiquitous, complex process fundamentally involved in the growth and proliferation of cells and entire organisms. The cell cycle represents an essential process governed by the coordination of highly controlled molecular processes. Numerous regulatory proteins and signalling pathways unite to direct the cell through a specific sequence of events. These result in the replication of the cell’s genetic material culminating at cell division, *i*.*e*. the event of equal partitioning of the cell’s genetic material and cell splitting into two newly-produced daughter cells (Schafer, 1998).

### Progression through cell cycle phases

The cell cycle can be sub-divided depending on various criteria; the most common categorisation groups the *G1-, S-*, and *G2-*phases, which together encompass the **interphase**. The interphase is the longest phase between subsequent cell divisions defined by cell growth and DNA replication event. Interphase represents the portion of the cell cycle that is not accompanied by visible cell changes under the microscope (Matthews et al., 2022), during which the cell partakes in various essential activities:

- In the **G1 (Gap / Growth one) phase**, the cell grows and responds to the environmental, biochemical and mechanical cues. A high amount of protein synthesis occurs and the cell accumulates nutrients and growth factors as it is about to double its original size. More organelles are produced and the volume of the cytoplasm increases.
- In the **Synthesis (S) phase**, the cell synthesises its DNA and the amount of DNA is doubled whilst the number of chromosomes remains constant, via semi-conservative replication (Hanawalt, 2004). Concomitantly, some of the organelles, including mitochondria, are replicated.
- In the **G2 (Gap / Growth two) phase**, the cell resumes its growth in preparation for division. The mitochondria (and in plants, also the chloroplasts) divide and the cell continues to grow and accumulate nutrients needed until mitosis begins.
- In addition, some cells that do not divide often (*e*.*g*. hepatocytes, stem cells) or ever (*e*.*g*. terminally differentiated cells, including epidermis, neurons, or muscle cells), enter a stage called **G0 (Gap zero) phase**, which is separate from, or a preceding extension of, the interphase. G0 cells are not actively cycling, and are therefore considered to interrupt the interphase. In some circumstances, the G0-phase may remain as a distinct quiescent stage which occurs outside of the cell cycle. Alternatively, the G0-phase may end and be followed by the remaining stages of interphase (Matthews et al., 2022).

The periods of interphase that surround the S-phase have historically been named the “gaps” in the cell cycle to separate the stages between the two obvious landmarks; the DNA synthesis and mitosis (Schafer, 1998). However, these phases are key periods for cell cycle regulation and include the crucial decision to commit to DNA replication during G1, and to initiate the process that leads to chromosome segregation during G2 (Matthews et al., 2022). In addition, to ensure that each interphase sub-stage is correctly completed, each phase ends when a **cellular checkpoint** validates the accuracy of the stage’s completion before proceeding to the next. As a quality control mechanism, the cell can decide to undergo apoptosis, which represents a highly regulated cell signalling process of biochemical and/or mechanical nature leading to programmed cell death.

## B. Appendix Deep image feature extractors

### Latent feature extraction

To extract image-based features describing PCNA cell phase morphology using automated approaches, cell-centred image patches were cropped from the fluorescence channel microscopy frames using the tracking coordinates. To complement manual feature extraction, we intended to learn latent representations of the image patches with an autoencoder-driven approach.

### Training image dataset

To train the variational autoencoder-based models, 3 representative movies had *≈*50, 000 patches randomly extracted. Since not all images expressed the *PCNA-iRFP* channel in sufficient amounts, the PCNA-reconstructing models were thus trained on particularly noisy, but representative, dataset.

### B.1. Vector Quantized Variational AutoEncoder (VQ-VAE)

#### *VQ-VAE* network architecture

The *VQ-VAE* architecture (Oord et al., 2018) is formed of 3 parts: first, a convolutional *encoder* compresses an input image *x* into a lower-dimensional representation *z*_*e*_(*x*), which is then *quantised* (by a nearest neighbour look-up, or *L*_2_ norm) along the channel dimension to a “codebook”, *i*.*e*. an embedding space *E∈*ℝ^*K×D*^. Here, *K* is the size of the discrete latent space, *i*.*e*. the number of available embeddings *e* in the codebook dictionary, and *D* is the dimensionality of each latent embedding vector *e*_*i*_. The quantised discrete embeddings *z*_*q*_(*x*) are then passed through a convolutional *decoder* to produce the output image *x*^*′*^, which is the image reconstruction of the same dimensionality as the original input *x* (Fig.8).

**Figure 8.**
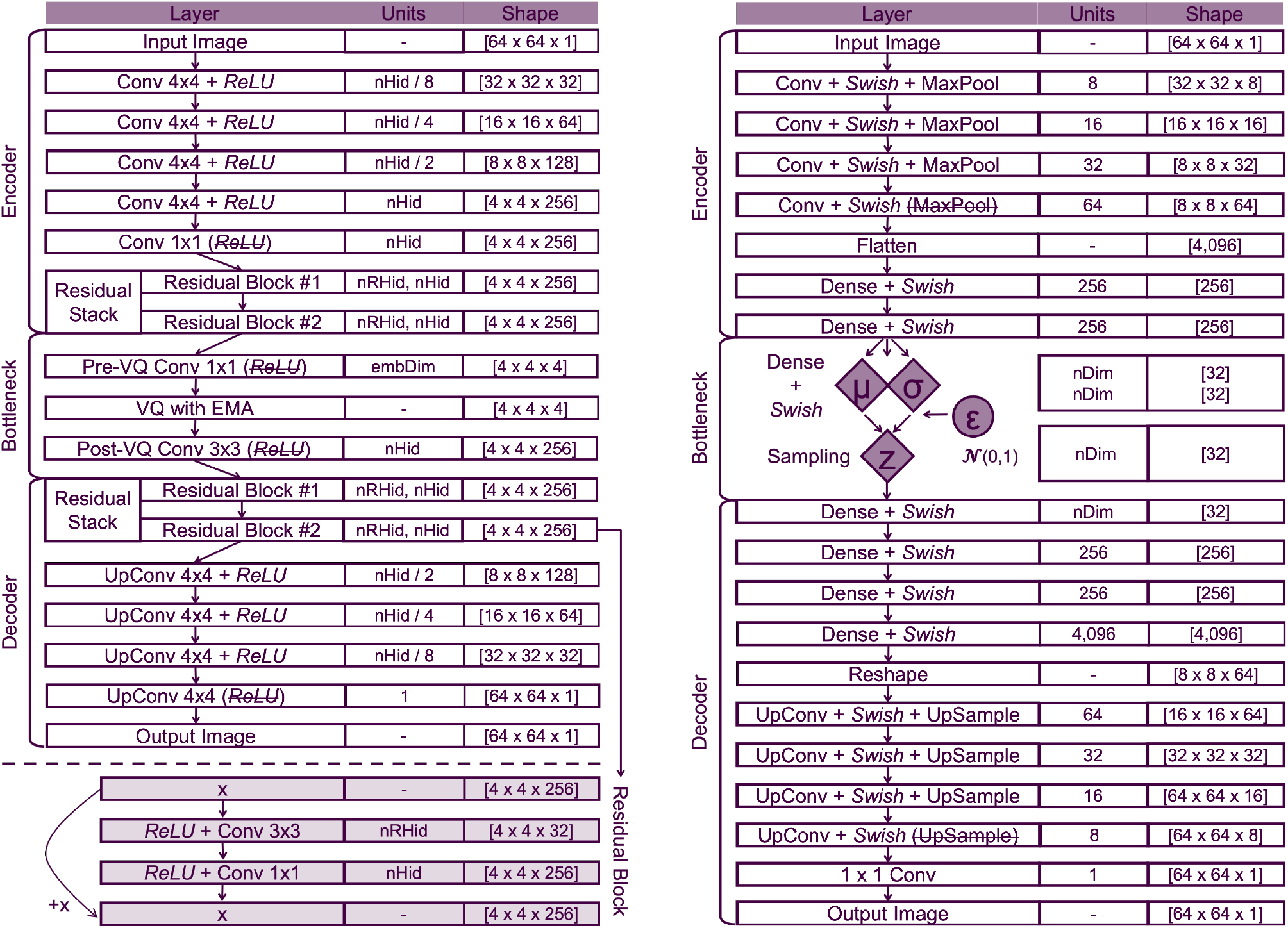
Architecture of the VQ-VAE (left) and *β*-VAE (right) models. *VQ-VAE* model architecture: The normalised single-channel input is funnelled through five convolutional layers, the output of which is flattened before being fed through a residual stack (illustrated at the bottom). The output of the residual stack is then fed into an additional convolutional layer to match the data dimensionality for the bottleneck where vector quantisation is performed. The entire process is reversed at the exit from the bottleneck and in the decoder step. to generate an output image with the identical dimensionality to the input image, [64, 64, 1]. Where (Up)Conv is specified, it suggests that an (up-)convolution 2D was performed with 2D *kernel_size* of 4*×*4 and stride of 2, or stride of 1 otherwise (kernel sizes 3*×*3 and 1*×*1). Rectified Linear Unit (ReLU) was used as an activation function throughout the architecture, and excluded at particular points to allow the downstream tensor to carry negative values (VQ-step, entry to residual stacks, output image). VQ, vector quantisation; EMA, exponential moving average; nHid, number of hidden units; nRHid, number of hidden units in the residual block; embDim, dimensionality of the embedding vector; +*x*, element-wise addition of original input (identity operation). *β****-VAE* model architecture:** The normalised single-channel input is funnelled through four convolutional layers, the output of which is flattened before being fed through two fully-connected (dense) layers. Then, the output is split into two separate dense layers, which represent the mean (*µ*) and log variance (*σ*) estimators of the distribution with specified dimensionality (here, 32). The *µ* and *σ* vectors are sampled from in the next layer using the Gaussian reparametrisation trick where *z* = *µ* + *σ ϵ* where *ϵ* 𝒩(*µ, σ*). This step separates the learnable (deterministic, diamond-shaped) parameters from the stochastic (circle) node to allow gradient flow during backpropagation. This randomly sampled information is then processed until the final layer, whose output is the same shape as the input. The final convolutional layer features 1*×*1 kernel that does not alter the image dimensions and omits the *Swish* activation function to allow the output to contain real negative pixel values. The decoder layers use nearest neighbour up-sampling between the convolutional layers to reverse the effect of the max-pooling layers in the encoder. For illustration, 32-dimensional space is used in the bottleneck step.

### Objective function

The *VQ-VAE* overall loss function is composed of 3 terms:

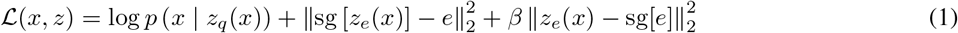

where the *reconstruction loss* (term 1) optimises both the encoder and the decoder of the model. In the forward pass, the nearest embedding *z*_*q*_(*x*) is passed to the decoder, but the codebook embeddings receive no gradients due to discretisation bottleneck bypassing in the backward pass. The *embedding loss* (term 2) is then minimised via *vector quantisation* based on *L*_2_ distance, to move the embedding vectors *e*_*i*_ towards the encoder outputs *z*_*e*_(*x*) by:

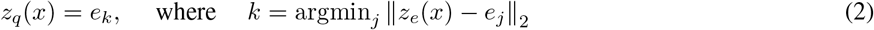

Whilst this term only updates the codebook dictionary, the *commitment loss* (term 3) of *z*_*e*_(*x*) to *e* prevents the dimensionless embedding space from growing arbitrarily. In addition, *β* = 0.25 is the scaling constant to balance the reconstruction and commitment losses, which was found, together with other hyperparameters such as letting the model train for 10,000 iterations with an embedding size = 256 and dimensionality = 4 via hyperparameter search.

### B.2. Disentangled Beta Variational AutoEncoder (*β*-VAE)

#### *β****-VAE* network architecture**

A convolutional variational autoencoder was built to learn a low-dimensional, interpretable representation of the cell image data, using the implementation from Soelistyo et al. (2022). The VAE is based on an encoder-decoder network crosstalk through a low-dimensional bottleneck - called the latent representation. The encoder part of the network is convolutional in nature, transforming the single-depth input patch **x** from the image space to the latent space. The dimensionality of the latent representation is specified in a separate hyperparameter. The decoder part of the network takes the latent mean of each feature in the representation and performs a series of reciprocal transformations to convert it back to an image space, yielding an reconstruction image, **x**^*′*^ (Fig.8).

The encoder network consisted of four convolutional layers with 3*×*3 kernels with 32, 64, 128 and 256 kernels, respectively, and Swish activation functions to introduce non-linearity. Each layer was pooled by a max-pooling operation with 2*×*2 pool size and stride. The convolutional output was flattened and passed through two FC layers with 256 units each – and swish activations - before being split into two branches - one FC layer branch was allowed to learn the mean, or *µ*, and the other one learned the variance, or *σ*^2^, estimator. For the Gaussian reparametrisarion trick, a random normal sampler was used to generate samples from the distribution. The architecture of the decoder is the inverse of that of the encoder, using nearest-neighbour upsampling between convolutional layers (Fig.8).

#### Objective function

The following objective function was used to train the network (Burgess et al., 2018; Higgins et al., 2017):

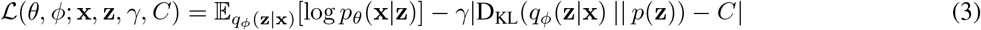

This loss function is composed of two competing terms. The combination of these two terms pushes the *β-VAE* to learn reliable representations of the image data, while regularising the latent space to promote a continuous representation of its latent space. Namely, the two components are:

1. The **reconstruction** component, which penalises differences between the original input image, **x**, and the reconstructed output image **x’**. The <u>m</u>ean squared <u>e</u>rror (MSE) function is implemented for the reconstruction loss.
2. The **regularisation** component, which implements the Kullback-Leibler divergence (D_KL_(*·* || *·*)). The aim of this is to penalise the latent space model variance from dropping to zero. This is done by forcing the encoding to match the prior distribution, here defined as a Gaussian prior with a diagonal covariance matrix, *N* (0, *I*).

The bottleneck capacity (*C*) of the network was dynamically adjusted during training. The value of *C* was scaled linearly as a function of training iteration, eventually reaching a maximum value *C*_max_ at 90% of training progression, remaining at the maximum plateau phase for the final epochs. This ensures that at early training iterations the network prioritise the encoding (through the regularisation loss), while at later iterations this is refined to optimise the decoding (through the reconstruction loss). The scaling constant *γ* balances the two terms of the loss function.

#### Pre-trained Residual Connection Network (ResNet)

The PyTorch implementation of the `ResNet-18` model (He et al., 2015) was used as an “off-the-shelf” feature extractor with default weights pre-trained on `ImageNet` dataset ^1^. The initial layer was modified to input a single-channel image and resize it to 224 *×* 224 pixel patch. The last layer was removed from the architecture, and the 512 features-long vector from the penultimate layer was used for the subsequent warping.

## C. Appendix: Dynamically warping time series latents

### Distance metric

An Euclidean (*ℓ*_2_) distance between two *L*-dimensional vectors (in this case, two image embeddings) is generalised as:

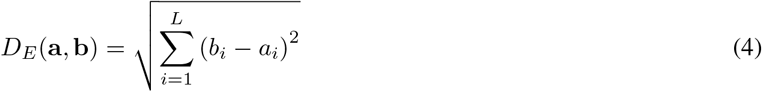

where a and b are the Cartesian vectors starting from the space origin, *a*_*i*_ and *b*_*i*_ are two points in Euclidean space where a, b *∈* R^*L*^.

### Dynamic time warping (DTW)

First, a pairwise cumulative warping matrix is computed (using Eq. 4) between any two trajectories. This matrix can be used to find the shortest DTW alignment path between the sequences of (non-)identical length using dynamic programming.

Two vectors *t* and *r* of lengths *m* and *n*, respectively, are found a mapping path *{*(*p*_1_, *q*_1_), (*p*_2_, *q*_2_), …, (*p*_*k*_, *q*_*k*_)*}* such that the distance on this mapping path 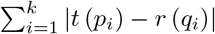 is minimized. In backward DTW, the optimum-value function is defined as *D*(*i, j*) as the DTW distance between *t*(*i* : *m*) and *r*(*j* : *n*), with the mapping path from (*i, j*) to (*m, n*). This recursively traverses the path as follows:

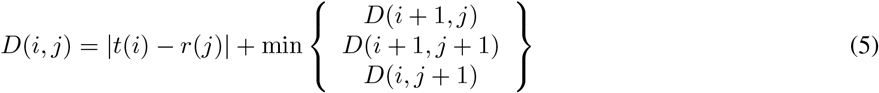

with the initial condition *D*(*m, n*) = |*t*(*m*) *− r*(*n*)| and the final answer of *D*(1, 1) (Berndt & Clifford, 1994).

## Footnotes

1 https://pytorch.org/vision/main/models/generated/torchvision.models.resnet18.html

## References

Berndt, D. J. and Clifford, J. Using dynamic time warping to find patterns in time series. In KDD Work-shop, 1994. URL https://dl.acm.org/doi/10.5555/3000850.3000887.

Bove, A., Gradeci, D., Fujita, Y., Banerjee, S., Char-ras, G., and Lowe, A. R. Local cellular neighborhood controls proliferation in cell competition. Molecular Biology of the Cell, 28(23):3215–3228, November 2017. ISSN 1059-1524, 1939-4586. doi: 10.1091/mbc.e17-06-0368. URL https://www.molbiolcell.org/doi/10.1091/mbc.e17-06-0368.

Burgess, C. P., Higgins, I., Pal, A., Matthey, L., Watters, N., Desjardins, G., and Lerchner, A. Understanding disentangling in β-vae, 2018. URL https://arxiv.org/abs/1804.03599.

El-Labban, A., Zisserman, A., Toyoda, Y., Bird, A., and Hyman, A. Temporal models for mitotic phase labelling. Medical Image Analysis, 18(7):977–988, 2014. ISSN 1361-8415. doi: https://doi.org/10.1016/j.media.2014.05.003. URL https://www.sciencedirect.com/science/article/pii/S1361841514000681.

Gallusser, B., Stieber, M., and Weigert, M. Self-supervised dense representation learning for live-cell microscopy with time arrow prediction, 2023. URL https://arxiv.org/abs/2305.05511.

Hanawalt, P. C. Density matters: The semiconservative replication of dna. Proceedings of the National Academy of Sciences, 101(52):17889–17894, 2004. doi: 10.1073/pnas.0407539101. URL https://www.pnas.org/doi/abs/10.1073/pnas.0407539101.

He, K., Zhang, X., Ren, S., and Sun, J. Deep residual learning for image recognition, 2015. URL https://arxiv.org/abs/1512.03385.

Higgins, I., Matthey, L., Pal, A., Burgess, C., Glorot, X., Botvinick, M., Mohamed, S., and Lerchner, A. beta-VAE: Learning basic visual concepts with a constrained variational framework. In International Conference on Learning Representations, 2017. URL https://openreview.net/forum?id=Sy2fzU9gl.

Ji, P., Zhang, T., Li, H., Salzmann, M., and Reid, I. Deep subspace clustering networks. Advances in Neural Information Processing Systems, 30, 2017. URL https://proceedings.neurips.cc/paper_files/paper/2017/file/e369853df766fa44e1ed0ff613f563bd-Paper.pdf.

Matthews, H. K., Bertoli, C., and de Bruin, R. A. M. Cell cycle control in cancer. Nature Reviews Molecular Cell Biology, 23(1):74–88, 2022. ISSN 1471-0080. doi: 10.1038/s41580-021-00404-3. URL https://doi.org/10.1038/s41580-021-00404-3.

Oord, A. v. d., Vinyals, O., and Kavukcuoglu, K. Neural Discrete Representation Learning, May 2018. URL http://arxiv.org/abs/1711.00937. arXiv:1711.00937 [cs].

Rappez, L., Rakhlin, A., Rigopoulos, A., Nikolenko, S., and Alexandrov, T. DeepCycle reconstructs a cyclic cell cycle trajectory from unsegmented cell images using convolutional neural networks. Molecular Systems Biology, 16(0), October 2020. ISSN 1744-4292, 1744-4292. doi: 10.15252/msb.20209474. URL https://onlinelibrary.wiley.com/doi/10.15252/msb.20209474.

Schafer, K. A. The cell cycle: A review. Veterinary Pathology, 35(6):461–478, 1998. doi: 10.1177/030098589803500601. URL https://doi.org/10.1177/030098589803500601. PMID: 9823588.

Shakarchy, A., Zarfati, G., Hazak, A., Mealem, R., Huk, K., Avinoam, O., and Zaritsky, A. Machine learning inference of continuous single-cell state transitions during myoblast differentiation and fusion. bioRxiv, 2023. doi: 10.1101/2023.02.19.529100.URL https://www.biorxiv.org/content/early/2023/02/19/2023.02.19.529100.

Soelistyo, C. J., Vallardi, G., Charras, G., and Lowe, A. R. Learning biophysical determinants of cell fate with deep neural networks. Nature Machine Intelligence, 4(7):636–644, June 2022. ISSN 2522-5839. doi: 10.1038/s42256-022-00503-6. URL https://www.nature.com/articles/s42256-022-00503-6.

Ulicna, K., Vallardi, G., Charras, G., and Lowe, A. R. Automated Deep Lineage Tree Analysis Using a Bayesian Single Cell Tracking Approach. Frontiers in Computer Science, 3:734559, October 2021. ISSN 2624-9898. doi: 10.3389/fcomp.2021.734559. URL https://www.frontiersin.org/articles/10.3389/fcomp.2021.734559/full.

Ulicna, K., Ho, L. T. L., Soelistyo, C. J., Day, N. J., and Lowe, A. R. Convolutional neural networks for classifying chromatin morphology in live-cell imaging. Chromosome Architecture: Methods and Protocols, pp. 17–30, 2022. doi: 10.1007/978-1-0716-2221-63. URL https://doi.org/10.1007/978-1-0716-2221-6_3.

Weigert, M., Schmidt, U., Haase, R., Sugawara, K., and Myers, G. Star-convex polyhedra for 3d object detection and segmentation in microscopy. In 2020 IEEE Winter Conference on Applications of Computer Vision (WACV). IEEE, mar 2020. doi: 10.1109/wacv45572.2020.9093435. URL https://www.computer.org/csdl/proceedings-article/wacv/2020/09093435/1jPbBJZQVZ6.

Wu, Z., Chhun, B. B., Popova, G., Guo, S.-M., Kim, C. N., Yeh, L.-H., Nowakowski, T., Zou, J., and Mehta, S. B. DynaMorph: self-supervised learning of morphodynamic states of live cells. Molecular Biology of the Cell, 33(6):ar59, May 2022. ISSN 1059-1524, 1939-4586. doi: 10.1091/mbc.E21-11-0561. URL https://www.molbiolcell.org/doi/10.1091/mbc.E21-11-0561.

